# Mathematical bounds on *r*^2^ and the effect size in case-control genome-wide association studies

**DOI:** 10.1101/2024.12.17.628943

**Authors:** Sanjana M. Paye, Michael D. Edge

## Abstract

Case-control genome-wide association studies (GWAS) are often used to find associations between genetic variants and diseases. When case-control GWAS are conducted, researchers must make decisions regarding how many cases and how many controls to include in the study. Depending on differing availability and cost of controls and cases, varying case fractions are used in case-control GWAS. Connections between variants and diseases are made using association statistics, including *χ*^2^. Previous work in population genetics has shown that LD statistics, including *r*^2^, are bounded by the allele frequencies in the population being studied. Since varying the case fraction changes sample allele frequencies, we extend use the known bounds on *r*^2^ to explore how variation in the fraction of cases included in a study can impact statistical power to detect associations. We analyze a simple mathematical model and use simulations to study a quantity proportional to the *χ*^2^ noncentrality parameter, which is closely related to *r*^2^, under various conditions. Varying the case fraction changes the *χ*^2^ noncentrality parameter, and by extension the statistical power, with effects depending on the dominance, penetrance, and frequency of the risk allele. Our framework explains previously observed results, such as asymmetries in power to detect risk vs. protective alleles, and the fact that a balanced sample of cases and controls does not always give the best power to detect associations, particularly for highly penetrant minor risk alleles that are either dominant or recessive. We show by simulation that our results can be used as a rough guide to statistical power for association tests other than *χ*^2^ tests of independence.

## Introduction

When conducting a genome-wide association study (GWAS), researchers search for trait-associated variants across an organism’s genome (Ikegawa 2012; Visscher, Wray, et al. 2017; Uffelmann et al. 2021). GWAS are often conducted for binary traits, in which the dependent variable expresses whether an individual has a trait of interest, such as a disease (Ozaki et al. 2002; Tanaka et al. 2003; Zondervan and Cardon 2004; Mototani et al. 2005). If the phenotype is a disease, study participants with the disease are called “cases,” and participants without the disease are “controls.” Case-control studies are common across epidemiology and related fields, where they are used to study potential risk factors for diseases by comparing their frequency in cases with their frequency in controls (Breslow 1996; DiPietro 2010). In a case-control GWAS, the putative risk factors are genotypes or alleles, and the signal of association is a difference in genotype or allele frequency between cases and controls.

To carry out a case-control study, one must decide the composition of the study sample. One key decision is setting the relative size of the samples of cases and controls, or the case fraction (Dupepe et al. 2019). The case fraction may affect statistical power to detect a risk factor in a case-control study. From first principles, with no information about the frequency of a putative risk factor in either cases or controls (and no difference in the cost of gathering data from cases vs. controls), a 1:1 ratio of cases and controls might be preferred: conditional on a given total sample size, a 1:1 ratio minimizes the standard error of the estimated difference in the frequency of the putative risk factor between cases and controls under the null hypothesis that the risk factor is at equal frequency in the two groups.^1^

Several researchers have considered the situation in more detail, motivated by differences in the difficulty or cost of collecting data from cases vs. controls (Ury 1975; Hennessy et al. 1999; Hong and Park 2012; Li et al. 2019). For many diseases, it is easier to recruit controls than cases, meaning that designs with more controls than cases are of interest (Dai et al. 2021). A common framework for planning matched case-control studies is to treat the number of cases as fixed and to examine how the study’s power changes as the number of matched controls per case increases (Gail et al. 1976; Ury 1975). In case-control GWAS, the rise of large biobank resources means that for any given disease, genetic data may be available from many people who might be considered for inclusion as controls. However, diseases that are rare in the general population will also likely be rare in a biobank, driving case fractions down well below 50%. This situation has motivated the development of new methods for GWAS that can accommodate extremely uneven samples of cases and controls (Zhou et al. 2018; Dai et al. 2021).

Another reason to consider the effect of varying the case fraction is that we may have some prior knowledge of the frequencies of risk or protective factors in the population. In particular, allele and genotype frequencies are subject to the evolutionary forces of drift, mutation, and selection. The balance of drift and mutation ensures that loci with low minor allele frequencies will outnumber those with higher minor allele frequencies, and for phenotype-associated variants, natural selection may also affect allele frequencies (Simons et al. 2022). Allele frequency affects statistical power in GWAS generally, and in case-control GWAS, it influences power in a way that depends on the case:control ratio. It has been observed that in case-control GWAS, there is often more power to detect loci with risk-increasing minor alleles than loci with protective minor alleles, particularly when considering loci with relatively large effects (Chan et al. 2014; Visscher, Hemani, et al. 2014).

Although in practice, many methods are used to analyze data in case-control GWAS, one way to approximate the power obtained in a case-control GWAS is by studying the non-centrality parameter governing the non-central *χ*^2^ distribution describing the distribution of the *χ*^2^ statistic from a test of independence between case status and genotype. The non-centrality parameter is closely related to the *r*^2^ measure of linkage disequilibrium (LD) used in population genetics. Specifically, for a haploid case-control GWAS, with a 2 × 2 table indicating the presence or absence of a putative risk allele on one dimension and case vs. control status on the other dimension, the noncentrality parameter is *nr*^2^, where *n* is the sample size and the *r*^2^ statistic is computed as if case vs. control status were a second “locus.” For *χ*^2^ tables with minimum dimension 2, as in case-control situations, the noncentrality parameter divided by *n* is equal to the square of Cramér’s *V*, a measure of effect size for associations between nominal variables.

Previous work in population genetics has explored bounds on statistics that are imposed by allele frequency in a population. The *r*^2^ statistic, in particular, is known to be bounded by the allele frequencies of the population being studied (VanLiere and Rosenberg 2008). This is one of many results in population genetics relating allele frequencies to mathematical bounds on statistics describing genetic diversity, LD, or population differentiation (Rosenberg and Jakobsson 2008; Jakobsson, Edge, and Rosenberg 2013; Edge and Rosenberg 2014; Alcala and Rosenberg 2016; Aw and Rosenberg 2018; Mehta et al. 2019; Kang and Rosenberg 2019; Alcala and Rosenberg 2022).

The relationship between the *χ*^2^ non-centrality parameter and LD statistics suggests that the non-centrality parameter is also bounded by allele frequencies in a case-control study. These bounds could explain observations about the power of case-control GWAS to detect the effects of different kinds of alleles, such as minor alleles that are risk-associated vs. protective (Chan et al. 2014; Visscher, Hemani, et al. 2014).

We analyze how varying the ratio of cases to controls in a case-control study affects the *χ*^2^ non-centrality parameter (Edwards et al. 2005; Visscher, Hemani, et al. 2014), adding to previous results by relating them to bounds on *r*^2^. We find that for variants with small effect sizes, the intuition underlying the 1:1 case-control ratio is justified. However, for large effect sizes, the bounds on the non-centrality parameter become important, and 1:1 case:control ratios become suboptimal. We use simulations to confirm that the intuition that comes from examining the bounds on *r*^2^ is a reasonable guide to the behavior of tests other than the Pearson *χ*^2^ test.

## Model

We consider a disease-associated biallelic locus in Hardy–Weinberg equilibrium. There are two possible alleles at the locus, denoted by *A* and *a*, with *a* being the disease-associated (“risk”) allele. We consider both a haploid case with two genotypes *A* and *a*, and a diploid case with three genotypes, *AA, Aa*, and *aa*. In both cases, we assume a binary disease phenotype.

Our notation is summarized in Table 1. The frequency of the disease allele in the population is represented by *p*. The frequency of disease cases in the population is denoted by *d*. The probability of having the disease given a genotype with no risk alleles is represented by *γ*.

**Table 1:**
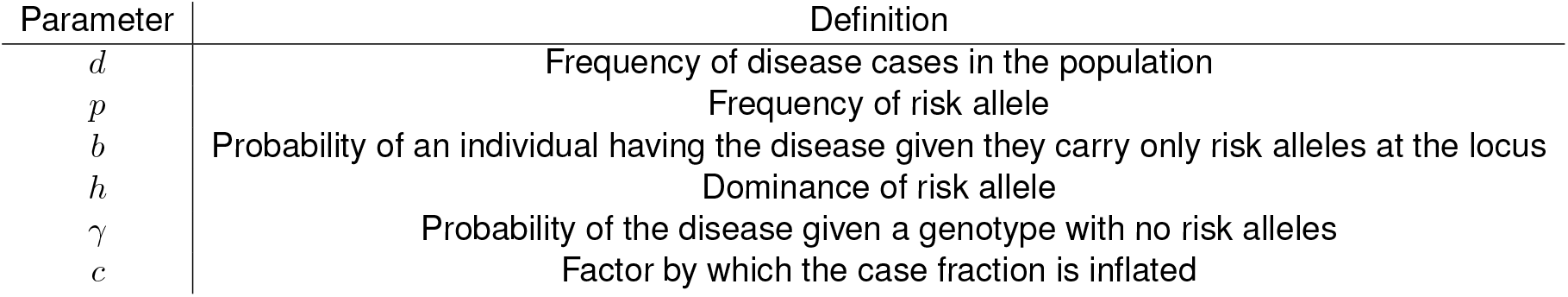
Summary of notation.

The effect size of the risk allele is governed by *b*, the penetrance, or the probability of developing the disease conditional on carrying only risk alleles at the locus. The penetrance *b* can, in principle, be less than *γ*, the disease risk for the protective genotype, but such values change the interpretation of the results (the “risk” allele becomes protective), so we focus on cases in which *b > γ*. In the diploid case, the dominance coefficient *h* controls whether the disease allele is dominant, recessive, or incompletely dominant. Specifically, the disease frequency among heterozygotes is *hb* + (1 − *h*)*γ*. When *h* = 1, the risk allele is fully dominant, and when *h* = 0, the risk allele is fully recessive. Although researchers sometimes assume an underlying normally distributed risk scale and define dominance with respect to this scale, we define dominance with respect to the probability of developing the disease. For any configuration of *γ, b*, and *h*, the same result could be obtained under a normal liability-threshold model with a different value of *h* chosen to give the same disease probabilities for heterozygotes as in our case.^2^ We also do not interpret values of *h* outside [0, 1], though much of our mathematical analysis applies to such cases.

In the haploid case, setting two values of *b, d*, and *γ* implies the value of the third, since *d* = *bp*+*γ*(1−*p*). In the diploid case, setting three of *b, d, γ*, and *h* implies the value of the fourth, since *d* = *bp*^2^ + [*hb* + (1 − *h*)*γ*]2*p*(1 − *p*) + *γ*(1 − *p*)^2^.

To allow for variation in the case fraction, we modified the frequency of the disease case cells by a factor *c*. That is, if the proportion of cases in the population is *d*, then the proportion of cases in the study sample is *cd*. Thus, *c* is the factor by which the number of cases is inflated in the study sample compared with the population at large. Because the proportion of cases in the sample must be less than 1, *c* is bounded from above; specifically, *c <* 1/*d*.

### *χ*^2^ effect size

Our main interest is in the effect size measuring departure from independence in the population contingency table relating genotype and disease status. The quantity we focus on, which we call *λ* and is sometimes called *ϕ*^2^ or *X*^2^ (Mirkin 2001), is equal to 1/*n* times the noncentrality parameter of the non-central *χ*^2^ distribution arising asymptotically from tests of independence of genotype and disease status, where *n* is the sample size. It is also equal to 1/*n* times the value of the *χ*^2^ statistic obtained from a sample with joint genotype and disease frequencies exactly matching those in the population. Specifically, consider a pair of nominal variables *X* ∈ {1, …, *k*_1_} and *Y* ∈ {1, …, *k*_2_}. Define the probability *P* (*X* = *i* ∩ *Y* = *j*) = *p*_*ij*_, and further define *P* (*X* = *i*) = *p*_*i*._ and *P* (*Y* = *j*) = *p*_.*j*_. Then the effect size is

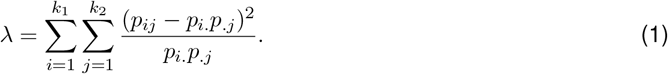

In our setting, one of the dimensions is a binary variable, case vs. control status, and the other is genotype. If we let *i* index genotypes, define *q*_*i*_ as the fraction of cases among individuals with genotype *i*, define *f*_*i*_ as the proportion of the sample with genotype *i*, and define *q* = Σ_*i*_ *f*_*i*_*q*_*i*_ the fraction of cases in the overall sample, then we can use the Brandt–Snedecor formula (Agresti 2013, p. 178) to write *λ* as

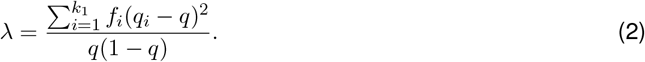

In this form, *λ* can be seen as a variance decomposition, which holds in more general *k*_1_ × *k*_2_ contingency tables (Mirkin 2001). If the fraction of cases in the sample is *q*, then the variance in case status for a random individual drawn from the sample is *q*(1 − *q*), and the between-genotype variance in the fraction of cases is the sum in the numerator. More specifically, if *D* is a random variable encoding case (*D* = 1) vs. control (*D* = 0) status, and *G* is a random variable encoding genotype, then equation 2 can be written as

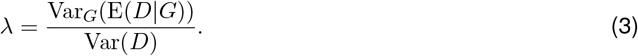

Equation 2 also allows us to express *λ* for a 2 × 3 contingency table as a weighted average of the *λ*s that emerge from the three possible 2 × 2 tables that result from omitting one of the columns. To start, note that the numerator can be re-expressed in terms of pairwise differences as follows, by remembering that *q* = Σ_*i*_ *f*_*i*_*q*_*i*_:

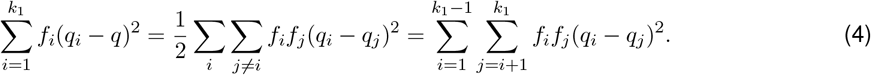

Next, define *λ*_*ij*_ as the value of *λ* that results from a 2 × 2 contingency table assembled from columns *i* and *j*,

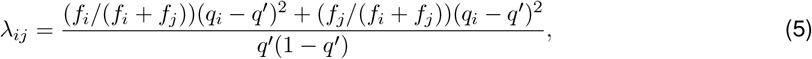

where *q*^*1*^ = (*q*_*i*_*f*_*i*_ + *q*_*j*_*f*_*j*_)/(*f*_*i*_ + *f*_*j*_). Using equation 4, we can write equation 5 as

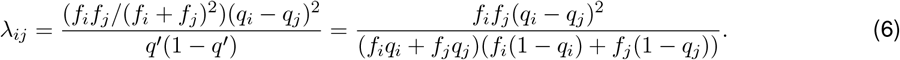

Combining equations 2, 4, and 6, we can re-express *λ* as a weighted sum of the *λ*_*ij*_ values,

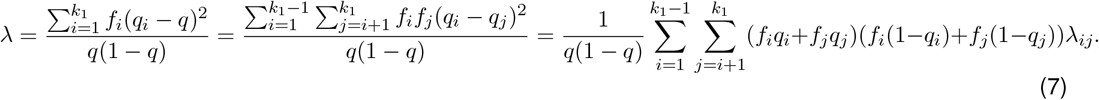

## Results

### Mathematical characterization of *λ*

#### Haploid case

The joint frequencies of disease and genotype (i.e. the *p*_*ij*_ terms in equation 1) in the haploid case are given in Table 2, along with the marginal frequencies (the *p*_*i*._ and *p*_.*j*_ terms in equation 1). To obtain these frequencies, start with the population frequencies (e.g. *P* (case ∩allele **a**) = *P* (case|allele **a**)*P* (allele **a**) = *bp*). Then multiply values in the case row by *c*, the factor by which case fraction in the sample differs from the population, and multiply values in the control row by (1 − *cd*)/(1 − *d*), the implied factor by which the control fraction in the sample differs from the population.

**Table 2:** Joint frequencies of a risk allele, **a**, a protective allele, **A**, and case vs. control status in a sample of haploids.

|  | **a** | **A** |  |
| --- | --- | --- | --- |
| **Controls** | $p(1 - b) \frac{1 - cd}{1 - d}$ | $(1 - d + bp) \frac{1 - cd}{1 - d}$ | $1 - cd$ |
| **Cases** | $cbp$ | $c(d - bp)$ | $cd$ |
| | $\frac{p(1 - cd - b + bc)}{1 - d}$ | $\frac{1 - d - p - pb(c - 1) + cdp}{1 - d}$ | |

Plugging these values into equation 1 gives *λ* in terms of the allele frequency *p*, the penetrance *b*, the overall disease frequency *d*, and the factor by which cases are oversampled compared with the population, *c* in the haploid case,

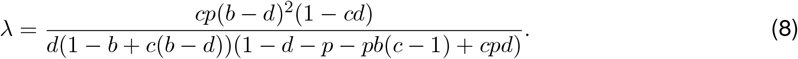

The expression for *λ* in equation 8 is closely related to the *r*^2^ measure of LD. In particular, it is equal to *r*^2^ if we think of case status and the risk allele as two “alleles” in LD in the sample. We can relate eq. 8 to the upper bounds on *r*^2^ in terms of allele frequency by considering a completely penetrant allele (i.e. *b* = 1). The upper bound on *r*^2^ takes different forms in each of eight triangles in the unit square describing the allele frequencies at the two loci under consideration (VanLiere and Rosenberg 2008). Since, for a completely penetrant risk allele, the disease frequency must be greater than or equal to the risk allele frequency, the corresponding bound on *r*^2^ in this case, if *p* and *d* are viewed as two allele frequencies, is

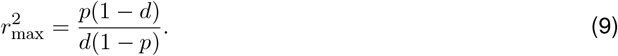

In our setting, disease frequency and allele frequency are modified from their population values by the parameter *c*. In particular, in the haploid case, the disease and allele frequencies in the sample can be expressed as *cd* and *cp*. With these sample frequencies, the function for the bound on *r*^2^ becomes

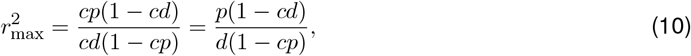

which is equivalent to the expression for *λ* in equation 8 when *b* is set to one. Therefore, in the haploid case, for a completely penetrant allele, the change in *λ* resulting from modifying the case fraction can be viewed as a traversal of the bounds on *r*^2^. In particular, changing the fraction of cases in the sample by modifying *c* is equivalent to traversing the surface that bounds *r*^2^ over a line that passes through the origin and the point (*d, p*).

Perhaps counterintuitively, for completely penetrant risk alleles, these paths along the surface imply that increasing the case fraction cannot increase the value of *λ*. The derivative of equation 10 with respect to the case sampling factor *c* is − *p*(*d* − *p*)/[*d*(1 − *cp*)^2^]. For the relevant setting (*p* ∈ (0, 1), *p* ∈ (0, *d*), *cp* ∈ (0, 1)), the derivative is negative unless the disease and risk allele frequency are equal (*d* = *p*), in which case it is zero (and *λ* = 1 for *cp* ≠ 1). (In our setting, *d* = *p* corresponds to a case in which the risk allele is both sufficient and necessary to develop the disease.)

An important caveat for interpreting this result in terms of statistical power is that the distribution of the *χ*^2^ statistic associated with the test of independence arising from this scenario has a noncentral *χ*^2^ distribution with noncentrality parameter equal to *nλ* only asymptotically. When some of the cells are empty, as is the case for a completely penetrant allele, the asymptotic distribution may not hold, and *λ* may not be a reliable guide to power. We explore this point by simulation later.

To consider a completely protective allele (*b* = 0), we can examine a region of the *r*^2^ bounds in which the disease frequency cannot be larger than one minus the protective allele frequency (*d* ≤ 1 − *p*), giving

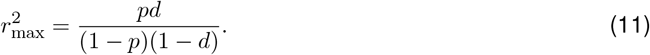

Setting the penetrance to *b* = 0 (i.e. the allele is completely protective) gives

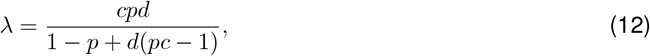

which is equal to equation 11 if *d* is set to *cd* and the frequency of the protective allele is set to *p*(1 − *cd*)/(1 − *d*), as would occur if cases are overrepresented in the sample compared with the population by a factor *c*. The derivative with respect to *c* of equation 12 is

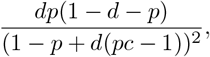

which, by the assumption that *d* ≤1 − *p*, is positive unless *d* = 1 − *p*, in which case it is zero (and *λ* = 1). Thus, for completely protective alleles, not surprisingly, the case is exactly reversed from that of a completely penetrant allele. The implication is that increasing the case fraction tends to increase *λ* for completely protective alleles, suggesting that power to detect protective vs. risk minor alleles will differ, and will respond to changes in the case fraction differently.

Therefore, in the haploid case, for both risk and protective alleles, when the allele’s effect is at maximum, the function for *λ* can be related to bounds on *r*^2^ (VanLiere & Rosenberg 2008). Varying the case fraction can be seen as moving along the surface of these bounds and changing the maximum value of *λ*, and thus the non-centrality parameter describing a *χ*^2^ test of independence applied to a case-control study (Figure 1).

**Figure 1:**
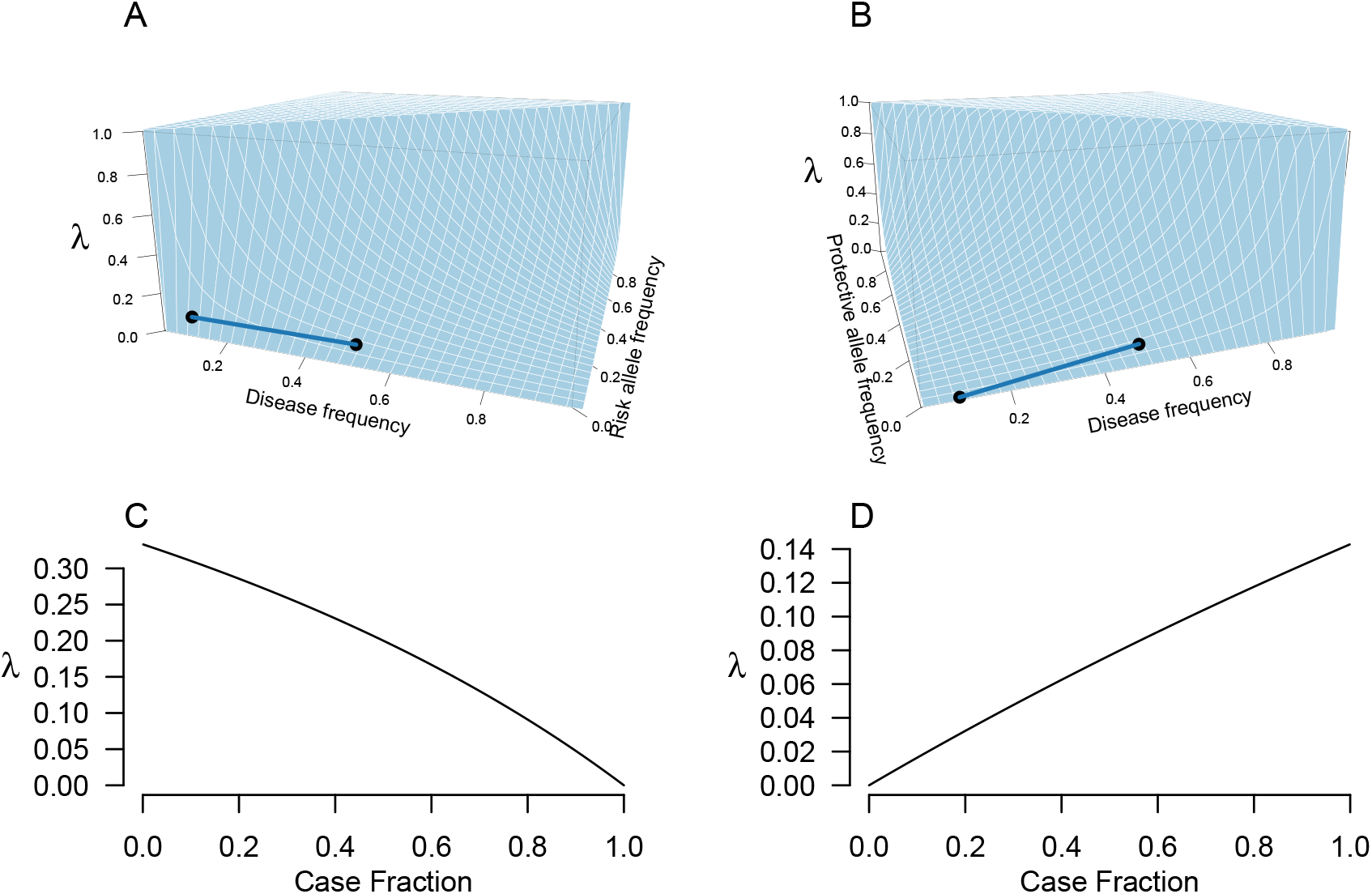
In the haploid case, when the penetrance *b* = 1, the change in the *χ*^2^ effect size *λ* that results from increasing the number of cases in the sample can be understood in terms of the bounds on the *r*^2^ LD statistic. A) The surface shows the value of *λ* as a function of the disease frequency *d* and the risk allele frequency *p*. The line connects the points on the surface immediately above (.1, .01) and (.01, .05), where *x* is the disease frequency and *y* is the frequency of the completely penetrant risk allele. The line represents the effect of increasing the percentage of cases in the sample from 10% to 50% and thereby increasing the frequency of the risk allele in the sample from 1% to 5%. B) Similar to (A), except that the *y* axis (into the page) now represents the frequency of a completely protective allele (*b* = 0). The line now represents changing the disease frequency in the sample from 10% to 50% and the protective allele frequency from 1% to 5%. C) A two-dimensional view of the traversal in (A) in terms of the fraction of cases in the sample. If an allele is completely penetrant but some individuals with the protective allele develop the disease, increasing the case fraction decreases *λ*. D) A two-dimensional view of the traversal in (B).

If we instead imagine an allele with a very small effect size, *λ* approaches a quadratic in *c*, the degree of case oversampling, maximized when the sample is evenly split between cases and controls. To see this, reparameterize equation 8 so that it is written in terms of Δ = *b* − *d*, the difference between the disease prevalence among carriers of the risk allele and the general population, rather than *b*. Doing so gives

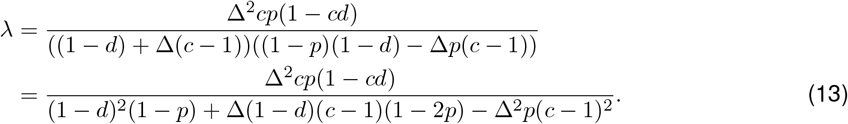

As Δ approaches 0 from above, the denominator of equation 13 is dominated by its first term, (1 − *d*)^2^(1 − *p*), which does not depend on *c*. Ignoring the other terms in the denominator makes equation 13 a concave quadratic in *c* with roots at 0 and 1/*d* (implying disease frequencies in the sample of 0 and 1) and a global maximum at *c* = 1/(2*d*) (implying a disease frequency in the sample of 1/2). Thus, we might expect that as the effect size of the risk variant decreases, *λ*’s dependence on the fraction of cases changes, such that for large effect sizes (i.e. near-complete penetrance), *λ* is maximized when the fraction of disease cases in the sample is close to the allele frequency in the sample, but for very small effect sizes (*b* − *d* ≈0), *λ* is maximized when the fraction of disease cases in the sample is approximately one half. This intuition matches our numerical results (Figure 2).

**Figure 2:**
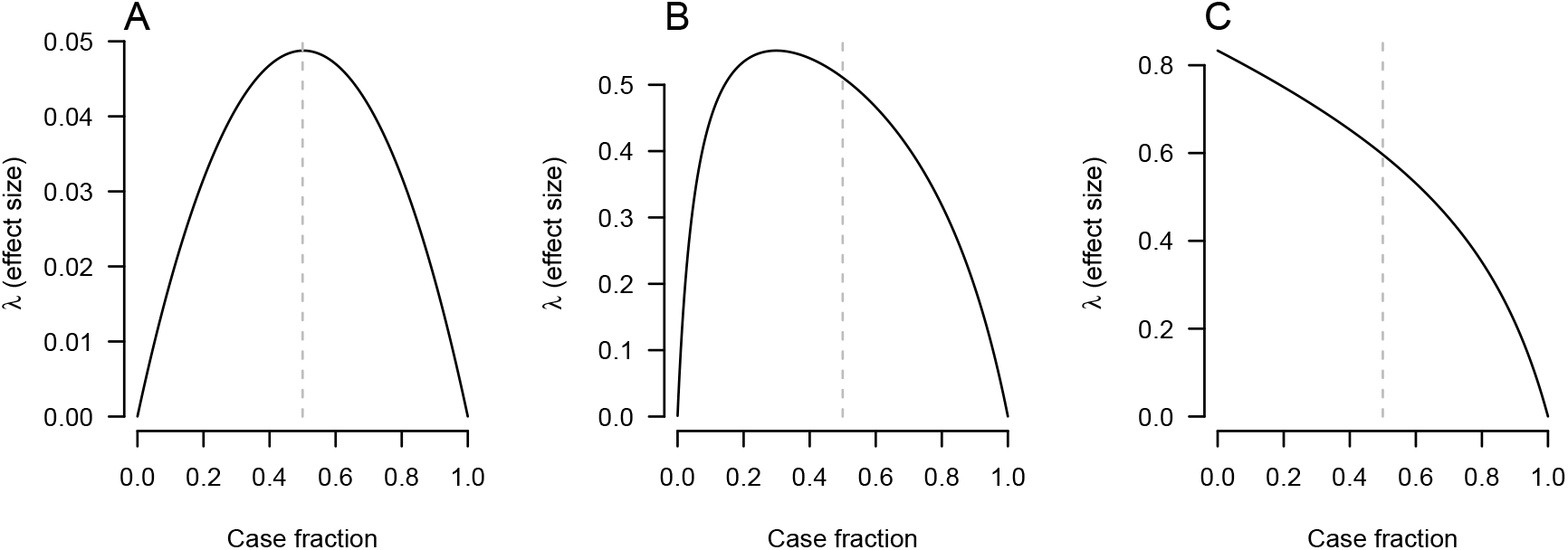
*λ* as a function of the case fraction in the haploid case at varying effect sizes. In all cases, the risk allele frequency *p* = 0.2 and the frequency of the disease among carriers of the protective allele is *γ* = 0.05. A) Penetrance *b* = 0.1, B) *b* = 0.8, C) *b* = 1. Vertical dashed lines indicate a sample with an equal number of cases and controls.

#### Diploid case

For diploids, we consider disease frequencies for three possible genotypes rather than two. The diploid case of the model extends the haploid case with the introduction of the dominance parameter, *h*, to specify the disease frequency for the heterozygous genotype. The joint frequencies for the three possible genotypes are shown in Table 3. In our parameterization, if the risk allele is dominant, then *h* = 1, and if the risk allele is recessive, then *h* = 0. If the risk allele is incompletely dominant, then *h* ∈ (0, 1).

**Table 3:** Joint frequencies of genotypes, **aa, Aa**, and **AA** case vs. control status in a sample of diploids. We assume that the locus is at Hardy–Weinberg equilibrium in the population.

|  | **aa** | **Aa** | **AA** |  |
| --- | --- | --- | --- | --- |
| **Controls** | $p^2(1-b)\frac{1-cd}{1-d}$ | $2p(1-p)(1-hb-\gamma(1-h))\frac{1-cd}{1-d}$ | $(1-p)^2(1-\gamma)\frac{1-cd}{1-d}$ | $1-cd$ |
| **Cases** | $p^2bc$ | $2p(1-p)(hb+\gamma(1-h))c$ | $(1-p)^2\gamma c$ | $cd$ |
| | $\frac{p^2(1-cd-b+bc)}{1-d}$ | $\frac{2p(1-p)(1-cd-\gamma+c\gamma-(1-c)(b-\gamma)h)}{1-d}$ | $\frac{(1-p)^2(1-c(d-\gamma)-\gamma)}{1-d}$ | |

The effect size *λ* can be written in terms of the parameters using equation 1 and the cells of table 2—internal cells correspond to the values of *p*_*ij*_, and the margins give the *p*_*i*._ and *p*_.*j*_ values. The resulting expression is unwieldy, but we can gain some insight into the effect of the bounds on *r*^2^ by recalling that *λ* in the diploid case can be expressed as a weighted sum of *λ* values from three different 2 × 2 contingency tables (equation 7). As such, *λ* is bounded by a function of the bounds on *r*^2^, namely a weighted average of the bounds computed for each of the three possible 2 × 2 tables formed from the columns of the 2 × 3 contingency table.

If one of the alleles is completely dominant (*h* = 0 or *h* = 1), then equations 2 and 4 reveal that *λ* is equal to the value it would take in a similar haploid situation. For concreteness, imagine that *h* = 0 and that the risk allele is therefore completely recessive. Let *q*_1_, *q*_2_, and *q*_3_ represent the fraction of cases among carriers in the sample of 0, 1, or 2 risk alleles, respectively. Then *h* = 0 implies that *q*_1_ = *q*_2_, and by equations 2 and 4,

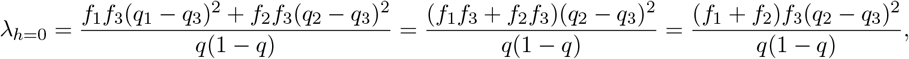

where the first simplification follows from applying the fact that *q*_1_ = *q*_2_. Thus, if the risk allele is fully recessive, then the effect size *λ* takes the value it would in a haploid scenario with the same *b* and *γ*, but protective allele frequency equal to the sum of the protective homozygote and heterozygote frequencies. By a similar argument, if the risk allele is fully dominant, then *λ* takes the value it would in an analogous haploid scenario, but with risk allele frequency equal to the sum of the risk homozygote and heterozygote frequencies. Thus, for fully recessive or dominant risk alleles, the arguments in the previous subsection apply directly.

Equation 7 reveals a second case in which the haploid results are straightforwardly applicable. If one of the alleles is rare, then one of the homozygotes will be very rare compared with the other genotypes. Thus, if the penetrance and case fraction are not too extreme, the weight (*f*_*i*_ in equation 7) on one of the homozygotes will be very small, causing it to contribute little to the value of *λ*. For example, for a risk allele at frequency 1% that is completely penetrant when homozygous and 50% penetrant in heterozygotes, there will be (1 − *p*)/*p* = 99 heterozygous cases for every homozygous case, causing risk homozygotes to contribute relatively little to *λ*, and implying that *λ* will be similar to the value of *λ* that would be obtained just by comparing heterozygotes with protective homozygotes.

For incomplete dominance and relatively common alleles, we find numerically that *λ* behaves broadly similarly to the haploid case, but with more of a tendency for case fractions near 1/2 to have relatively high *λ* values (Figure 3). Specifically, for low-penetrance alleles, *λ* looks like a concave quadratic in *c*, maximized when the fraction of cases in the sample is approximately 1/2. For higher-penetrance alleles and *d <* 1/2 (i.e. diseases at less than 50% frequency in the population), *λ* is maximized when disease frequencies in the sample are lower, closer to the population frequency. However, compared with the haploid case, the dependence of the sample case fraction that optimizes *λ* on penetrance is less for the diploid case, at least for intermediate values of the dominance parameter *h*.

**Figure 3:**
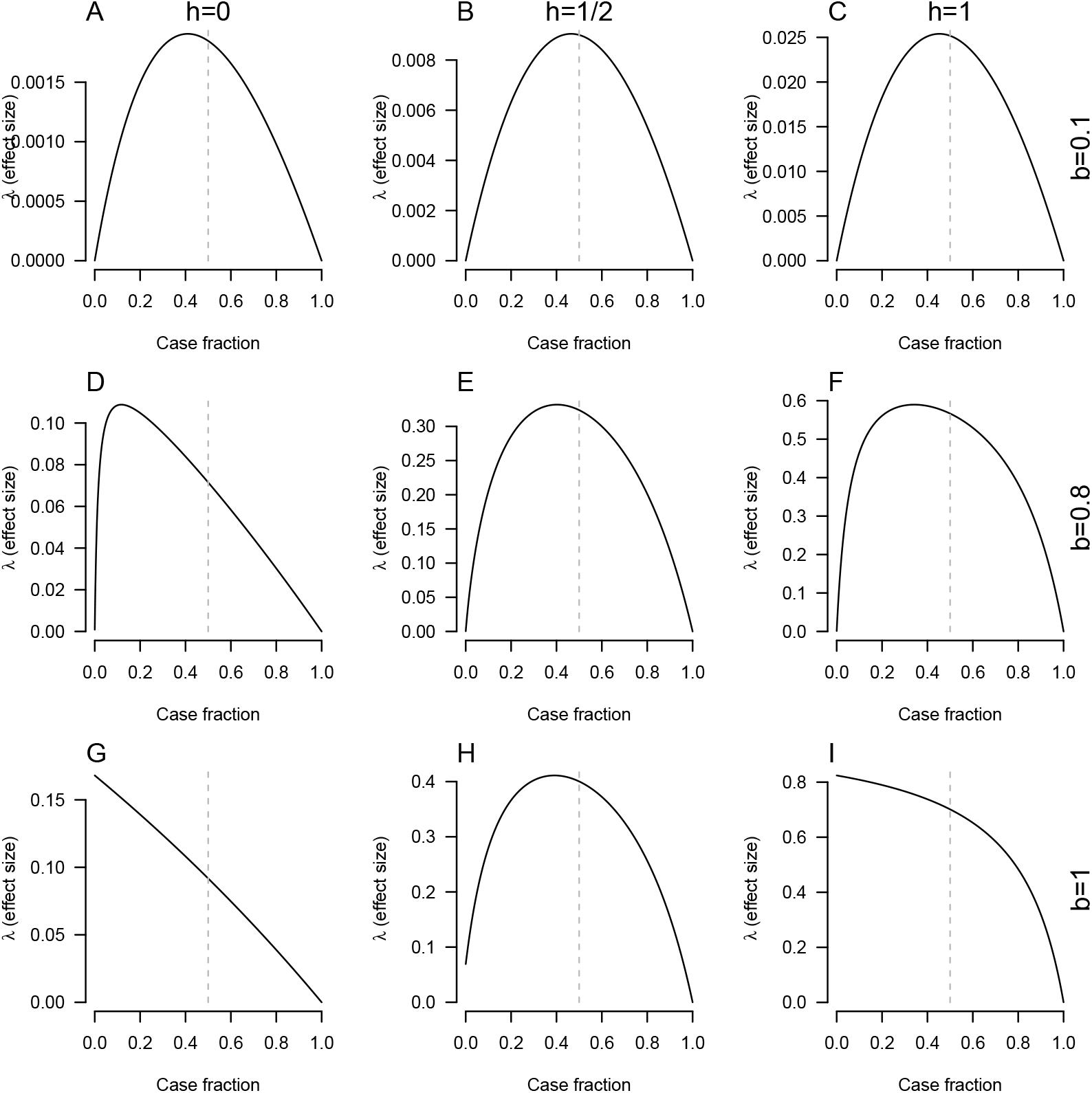
*λ* as a function of the case fraction in the diploid case. Each column displays a different dominance coefficient (*h* = 0, *h* = 1/2, and *h* = 1), and each row a different penetrance (*b* = 1/10, *b* = 4/5, and *b* = 1). In all panels, the risk allele frequency *p* = 1/10 and the probability of developing the disease among individuals with two copies of the protective allele is *γ* = 1/20. The vertical dashed grey lines indicate a case fraction of 1/2.

These observations can be understood in terms of the haploid results. When penetrance is low, the diploid *λ* can be seen as a weighted average of three haploid *λ*s, each of which has approximately the same shape—that of a concave quadratic function maximized when the disease fraction in the sample is 1/2.

Considering the high-penetrance case, with *b* = 1 and *h* = 1/2, *λ* becomes

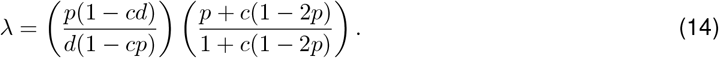

The first parenthetical term in the product in equation 14 is identical to equation 10, the haploid value of *λ* with complete penetrance, interpretable in terms of the bounds on the *r*^2^ LD statistic. As shown in the previous subsection, it is decreasing in *c* if *d > p* and *p >* 0. (It is guaranteed that *d* ≥ *p* if *h* = 1/2 and *b* = 1.) For allele frequencies *p <* 1/2, the second parenthetical term increases monotonically in *c*, equal to *p* when *c* = 0, to 1/2 when *c* = 1, and growing to 1 as *c* approaches infinity. (In our setting, *c* is bounded from above by 1/*d*.) Numerically, we observe that the second term acts to dampen the dependence of the the relationship between *λ* and *c* on the effect size, such that even for large effect sizes, if *h* = 1/2, then *λ* is maximized if the proportion of cases in the sample exceeds the proportion in the population (i.e. *c >* 1). Figure 4 shows additional diploid *λ* values, focusing on whether the minor allele is protective of risk-conveying.

**Figure 4:**
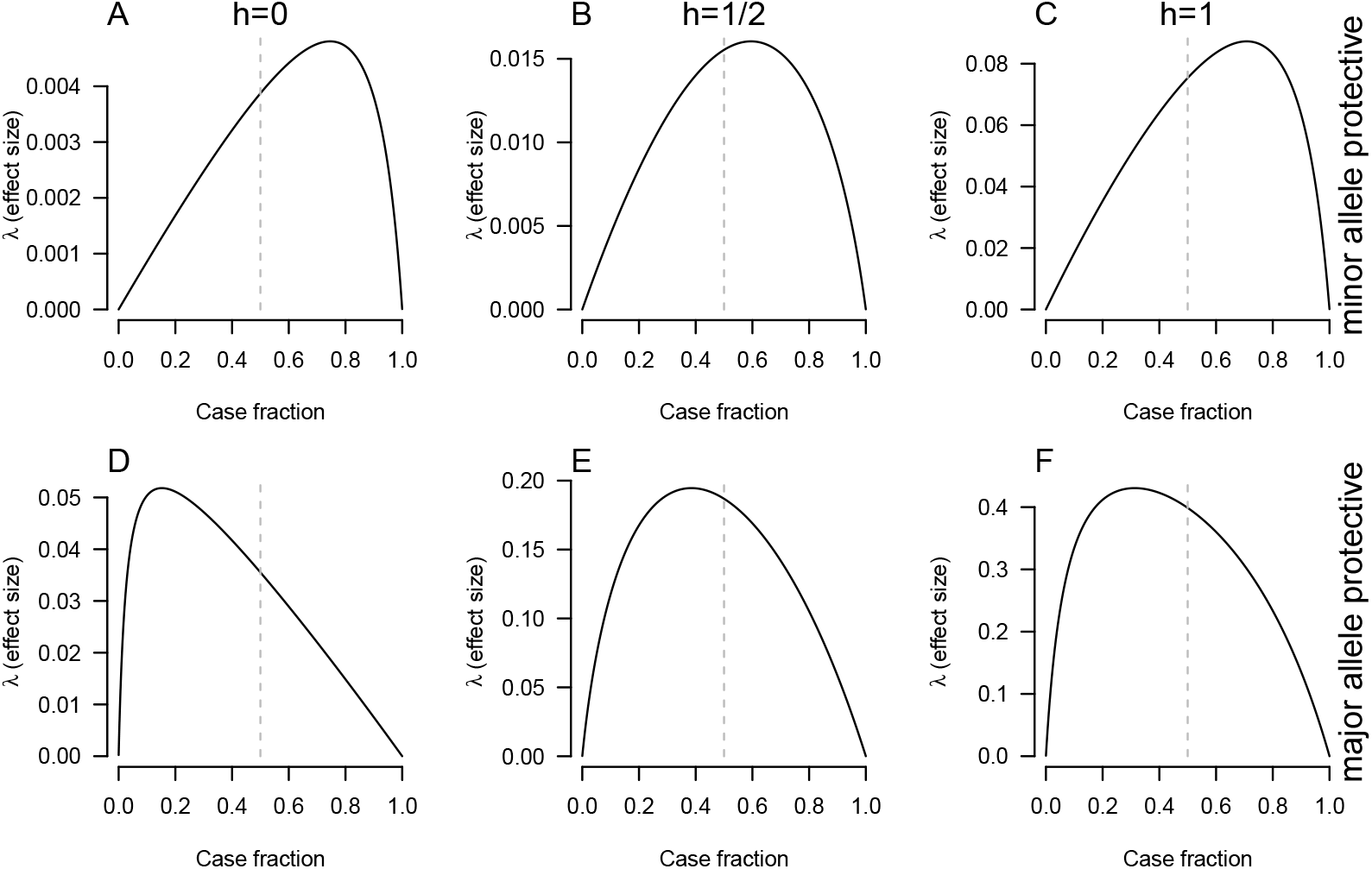
As in the haploid case, the highest value of *λ* as a function of the case fraction occurs when the case fraction is *>* 1/2 if the minor allele is protective, and when the case fraction is *<* 1/2 if the risk allele is the minor allele. In all panels, the minor allele frequency is *p* = 1/10 and the major allele homozygote has disease risk 1/10. In panels A-C, the minor allele homozygote has disease risk 1/80. In panels D-F, the minor allele homozygote has disease risk 4/5. In the left column, the minor allele is recessive; in the middle, the alleles are incompletely dominant (*h* = 1/2), and in the right column, the minor allele is dominant. The vertical dashed grey lines indicate a case fraction of 1/2.

### Diploid power simulations

Our mathematical results in the previous subsection describe the effect-size *λ*, which is proportional to the noncentrality parameter of the asymptotic distribution of the Pearson *χ*^2^ statistic computed from a contingency table of genotype vs. disease status. The noncentrality parameter determines the power of the test if the *χ*^2^ statisic indeed follows its asymptotic distribution. We investigated the degree to which our mathematical results are a valid guide to empirical power obtained in simulations.

We simulated genotype-by-case-status contingency tables obeying the probabilities in Table 3, fixing the row totals (i.e. forcing exactly the desired fraction of cases). We then computed Pearson *χ*^2^ tests on the resulting contingency tables and compared the fraction significant at level 5 × 10^−8^ with predictions obtained from the theoretical distribution.

Simulation results for a range of effect sizes and dominance coefficients are shown in Figure 5. For low-penetrance alleles, observed power is close to the predicted values. For higher-penetrance risk alleles, there are noticeable departures from theory, perhaps in part because simulated sample sizes are lower. (Sample sizes were chosen so that the maximum theoretical power value predicted from *λ* was approximately 0.9 in all cases.) However, the simulations support the qualitative predictions from the calculations, including that, for highly penetrant, recessive, minor risk alleles, power is optimized when the fraction of cases in the sample is substantially less than 1/2.

**Figure 5:**
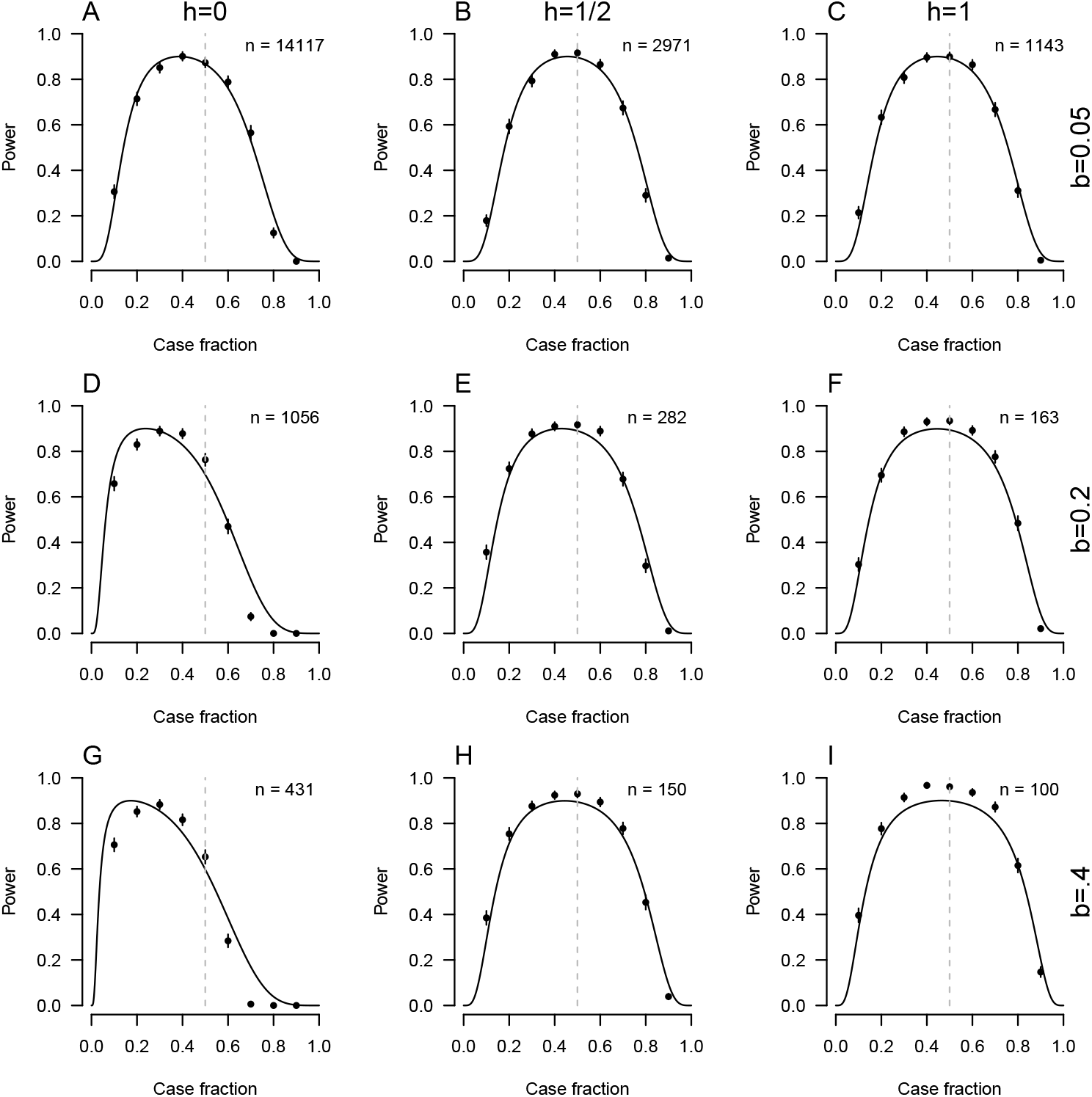
Predicted power (solid line) and empirical power from simulations (points) for Pearson’s *χ*^2^ tests of independence of diploid genotype and disease status. In all panels, the risk allele frequency *p* = 1/10, and the frequency of the disease among protective-allele homozygotes is *γ* = 1/50. Sample sizes were chosen to achieve a maximum predicted power of 90% and are printed in each panel. In panels A-C, the penetrance *b* = .05. In panels D-F, *b* = .2, and in G-I, *b* = .4. In panels A, D, and G, the risk allele is recessive (*h* = 0); in B, E, H, the risk allele is additive (*h* = 1/2), and in C, F, I, the risk allele is dominant (*h* = 1). Error bars on empirical power estimates represent *±*2 standard errors.

From the results of Figure 5, it appears an especially interesting case is that of a fully recessive, highly penetrant risk allele. We consider more examples of such alleles in Figure 6. In this case, the optimal case fraction is less than 1/2, and a sample with 1/2 cases has substantially lower power than samples with balanced cases and controls. Because fully recessive and fully dominant alleles can both be related exactly to the haploid case, equivalent results could be obtained with dominant risk alleles at lower frequencies.

**Figure 6:**
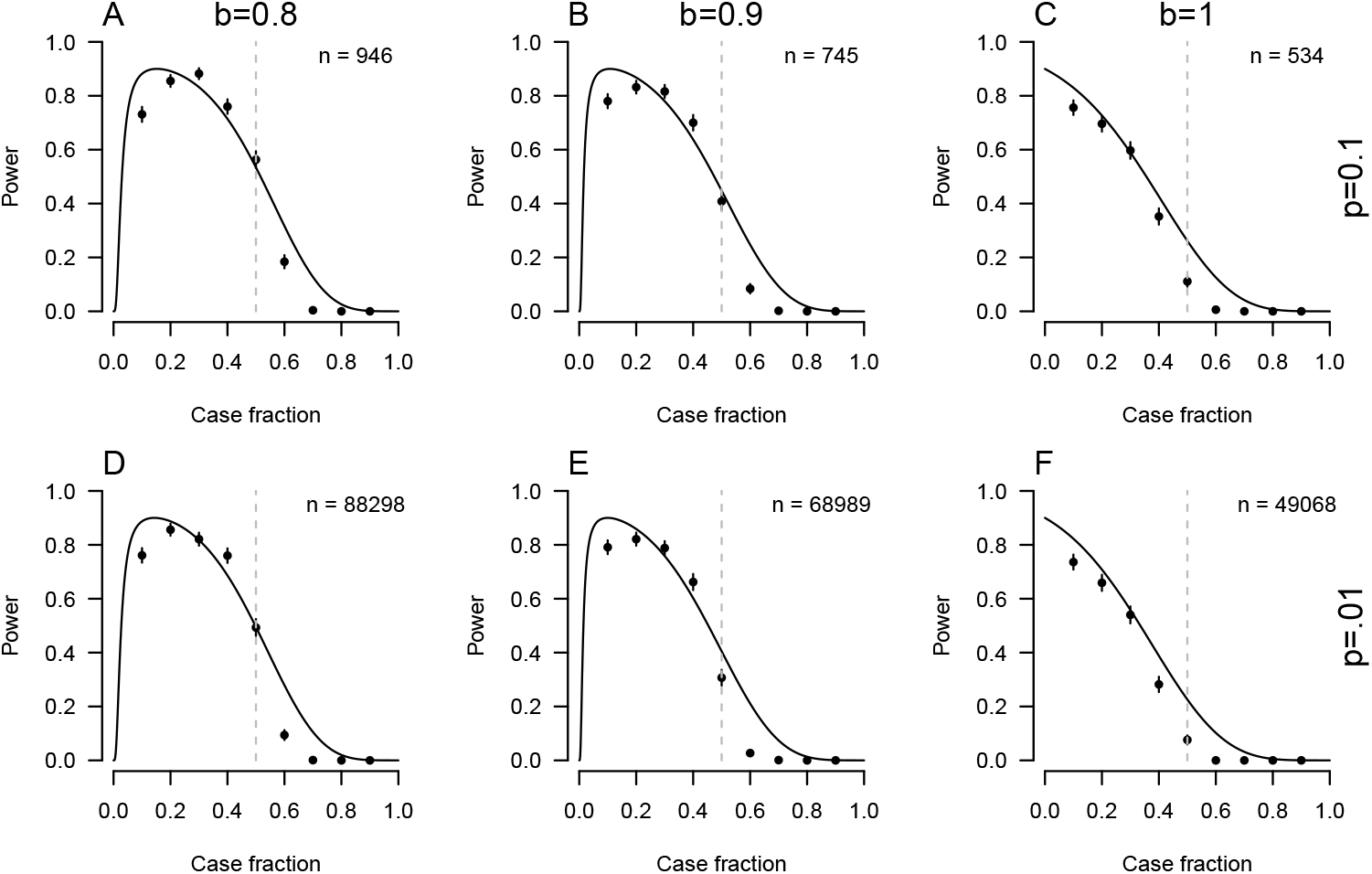
Predicted and empirical power for highly penetrant, fully recessive (*h* = 0) risk alleles. Conventions are as in Figure 5. In all panels, the disease risk among protective-allele homozygotes is *γ* = 1/10. In panels A-C, the risk allele frequency is *p* = 1/10, and in panels B-D, *p* = 1/100. From left to right, penetrance increases: *b* = 4/5 in the left column, *b* = 9/10 in the middle column, and *b* = 1 on the right.

### Other statistical tests

We have focused on the Pearson *χ*^2^ test for independence because it is a natural way to test for associations between genotype and a categorical outcome, and because it can be related to the *r*^2^ measure of LD and its known bounds, as we have shown. However, in practice, other methods are often used to test for associations between genotype and case status. In particular, researchers often use the Cochran– Armitage trend test (Cochran 1954; Armitage 1955) or a generalized linear model. The trend test often has an advantage of higher power when risk alleles act additively, and generalized linear models offer natural ways to adjust for covariates.

Figure 7 shows simulation results analogous similar to those in Figures 5 and 6, but including additional tests—a trend test and two generalized linear models, logisitic regression and probit regression. As in Figures 5 and 6, the Pearson *χ*^2^ test performs roughly as expected, with some noticeable deviations in the more extreme scenarios. As expected, the trend test usually outperforms the Pearson *χ*^2^ test when the risk allele is additive and underperforms when it is fully recessive. The generalized linear models struggle in some of the scenarios simulated here but perform similarly to the *χ*^2^ test in the case closest to their intended use (moderate effect size, additive risk allele). Notably, the other tests tend to follow the broad patterns predicted on the basis of *λ*, in particular higher power when the fraction of cases is below 1/2 for highly penetrant minor risk alleles.

**Figure 7:**
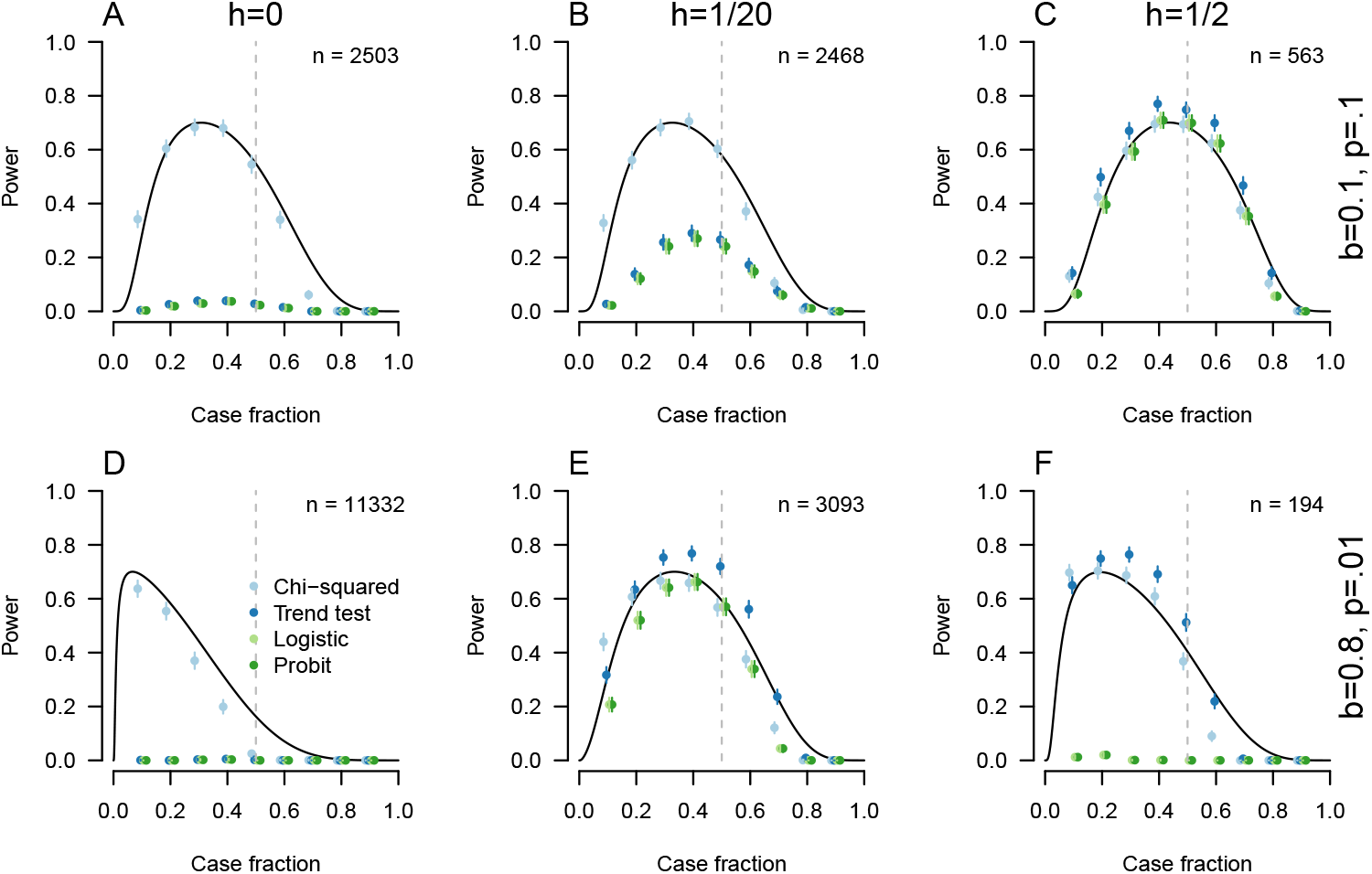
Predicted power for the Pearson *χ*^2^ test, and empirical power estimates from simulations for the *χ*^2^ test, Cochran–Armitage trend test, logistic regression, and probit regression. In all panels, the frequency of the disease among protective-allele homozygotes is *γ* = 1/50. Sample sizes were chosen to achieve a maximum predicted power for the *χ*^2^ test of 70% and are printed in each panel. In panels A-C, the risk allele is moderately penetrant (*b* = 1/10) and somewhat common (*p* = 1/10). In panels D-F, the risk allele is highly penetrant (*b* = 4/5) and rarer (*p* = 1/100). In the leftmost column, the risk allele is completely recessive (*h* = 0). In the middle column, it is not completely recessive (*h* = 1/20), and in the right column, it is additive (*h* = 1/2).

## Discussion

Motivated by the relationship between *r*^2^ measure of linkage disequilibrium and the non-centrality parameter arising from a *χ*^2^ test of independence in case-control GWAS, we have examined how variation in the fraction of cases used in a case-control study affects power to detect associations between genetic variants and diseases. The bounds on *r*^2^ in terms of the allele frequencies of the loci whose LD is being characterized (VanLiere and Rosenberg 2008) also characterize the value of the *χ*^2^ effect size *λ* for a completely penetrant risk allele in a haploid case-control GWAS. Varying the case fraction can be seen as moving *λ* along these bounds. For diploids, the haploid results can be applied directly if the risk allele is completely dominant or recessive, and they can be used to understand some cases with incomplete dominance as well, though such cases sometimes become unwieldy. Simulations support our approach as a means to understanding power in case-control GWAS, even with tests other than the Pearson *χ*^2^.

Depending on the dominance, penetrance, and frequency of the allele being studied, as well as the risk for the disease among individuals without the risk allele, the optimal case fraction for a fixed total sample size varies. Case fractions close to 50% are best for weakly penetrant risk alleles. As the penetrance of the risk allele increases, then for minor risk alleles, lower case fractions are expected to increase power, as the case fraction that maximizes *λ* decreases. Simulations support this assertion in the diploid case, though the effect is often small unless the allele is close to fully recessive (or dominant), in which case it can be quite pronounced.

In humans, massive datasets and other resources already exist for GWAS (Visscher, Wray, et al. 2017), and it is likely that the great majority of common, highly penetrant risk alleles have been found for well-studied diseases. Thus, in humans, it is likely that the results here are most practically useful for thinking about either low-penetrance alleles—in which case the intuition of attempting to balance cases and controls (given a fixed total sample size) is supported—or for considering the design of emerging sequencing studies of rare disease (Investigators 2021).

Several considerations left out of our model will also be important when considering such design choices (or indeed, in other organisms in which GWAS resources are not as developed). First, we do not consider the difference in cost of recruiting cases and controls. We instead consider the effect of varying the fraction of cases given a fixed total sample size. For rare diseases, it may be much easier to locate controls than cases. And in fact, large datasets of potential controls are generally widely available, depending on the epidemiological principles on which controls are selected. This will tend to push the optimal fraction of cases down, since many controls might be gathered for the cost of a single case. Our results suggest that this situation will make minor risk alleles easier to detect than minor protective alleles, an asymmetry that has been noticed before (Chan et al. 2014).

Second, we do not explicitly consider the possibility that we may test a marker allele rather than the causal allele itself. For a test at a non-causal marker, the *r*^2^-sense LD between the marker and the underlying causal allele(s) influences the power of the test (Pritchard and Przeworski 2001; Zondervan and Cardon 2004; Edge, Gorroochurn, and Rosenberg 2013). Thus, the bounds on *r*^2^ may need to be considered both with respect to the similarity in frequency of the causal and marker alleles (VanLiere and Rosenberg 2008) and with respect to the frequency of cases in the sample, as explored here. Allelic heterogeneity may also be prevalent in genes carrying highly penetrant risk alleles (Terwilliger and Weiss 1998), and such allelic heterogeneity may be better handled by approaches other than GWAS (Browning and Thompson 2012; Link et al. 2023).

Third, our model considers power to detect risk loci given a fixed allele frequency, dominance, effect size, and disease frequency. In practice, the allele frequencies and effect sizes of causal variants are not known, but it may be possible to develop predictions for effect size and allele frequency given parameters governing evolution of trait-associated loci, or to estimate aspects of the genetic architecture via other means. Integrating our functions over such joint distributions could provide guidance about case-control study design. Rough knowledge of genetic architecture also influences other aspects of study design, such as whether to focus on recruitment of cases with family histories of disease (Antoniou and Easton 2003; Zondervan and Cardon 2007).

Many important statistics in genetics are functions of allele frequencies, meaning that their arguments are non-negative and sum to one. The effects of such constraints have been explored in some detail in population genetics—they often lead to mathematical bounds that can explain counterintuitive aspects of the behavior of population-genetic statistics (Rosenberg and Jakobsson 2008; Jakobsson, Edge, and Rosenberg 2013; Edge and Rosenberg 2014; Alcala and Rosenberg 2016; Aw and Rosenberg 2018; Mehta et al. 2019; Kang and Rosenberg 2019; Alcala and Rosenberg 2022). These arguments have implications in other fields that use analogous statistics (Rosenberg and Zulman 2020), including in statistical genetics and genetic epidemiology.

## Acknowledgments

We thank members of the Edge, Mooney, and Pennell labs for helpful discussions. Funding was provided by NIH grant R35GM137758 to MDE.

## Code Availability

R code to produce all figures included here is available at https://github.com/mdedge/casecontrolandr2. All figures were produced using R version 4.1.2.

## Footnotes

1 To see this, let *p*_0_ and *p*_1_ be the true frequencies of the risk factor in controls and cases, respectively, and let *n*_0_ and *n*_1_ be the sample sizes of controls and cases, with *n* = *n*_0_ + *n*_1_ fixed. Assuming the control and case samples are independent, the variance of the difference in sample frequencies is 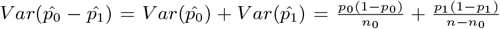. To minimize in terms of *n*_0_, we take the derivative to get 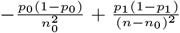. Recalling *n* − *n*_0_ = *n*_1_ and setting to zero gives an optimum where 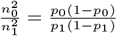, which is satisfied by setting *n* = *n* if the null hypothesis is true and *p*_0_ = *p*_1_.

2 Specifically, for a standard-normal liability-threshold model, our choice of *γ* implies a standard-normal liability of *γ*, = Φ^−1^(*γ*) for individuals with no risk alleles, where Φ is the cumulative distribution function of the standard normal. The penetrance *b*, similarly implies a normal liability *b*, = Φ^−1^(*b*) for individuals carrying only risk alleles. The dominance on the normal liability scale, *h*,, that corresponds to our choice of dominance coefficient *h*, is the solution of *h*,*b*, + (1 − *h*,)*γ*, = Φ^−1^(*hb* + (1 − *h*)*γ*), Which is 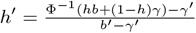.

## Notes

https://github.com/mdedge/casecontrolandr2

### Competing Interest Statement

The authors have declared no competing interest.

